# Unpacking sublinear growth: diversity, stability and coexistence

**DOI:** 10.1101/2024.06.03.597151

**Authors:** Guim Aguadé-Gorgorió, Ismaël Lajaaiti, Jean-François Arnoldi, Sonia Kéfi

## Abstract

How can many species coexist in natural ecosystems remains a fundamental question in ecology. Theory suggests that competition for space and resources should maintain the number of coexisting species far below the staggering diversity commonly found in nature. A recent model finds that, when sublinear growth rates of species are coupled with competition, species diversity can stabilize community dynamics. This, in turn, is suggested to explain the coexistence of many species in natural ecosystems. In this brief note we clarify why the sublinear growth (SG) model does not solve the long standing paradox of species coexistence. This is because in the SG model coexistence emerges from an unrealistic property, in which species per-capita growth rate diverges at low abundance, preventing species from ever going extinct. When infinite growth at low abundance is reconciled with more realistic assumptions, the SG model recovers the expected paradox: increasing diversity leads to competitive exclusion and species extinctions.

---

Naturalists in the 19th century had already realized that species competition, encapsulated in the notion of the *survival of the fittest*, was at odds with the diversity observed in natural ecosystems. If species compete for space or resources, one would not expect to find the high levels of diversity that are observed in *e*.*g* coral reefs or tropical forests [1–3]. Later, Robert May investigated this mathematically and proposed that a large, randomly interacting community becomes linearly unstable if diversity overcomes a predictable threshold [4]. More recent studies have further explored the dynamics of complex ecological models with competitive interactions, showing that, as diversity increases, species go extinct and fluctuations emerge [5–7].

Many solutions to that paradox have been suggested in the literature. Among others, natural communities are not random: the distribution of interaction strengths [8–10] or the structure of ecological networks [11–13] can increase the probability that many species coexist. These mechanisms allow communities to support a higher number of species than what would be expected if interactions strengths are random. However, they do not invert the shape of the dependency: a high diversity will still be detrimental for coexistence (see Box).

A recent study finds that a sublinear growth (SG) rate coupled with bilinear competition between species can lead to a reverse relationship, in which diversity begets stability [14]. More diverse communities would then be more resilient to perturbations in species abundances (see Box). This, in turn, is used to suggest that diversity can prevent competitive exclusion. If this was so, the model would propose a direct solution to the paradox of species coexistence under competition [2, 3].

The study presents strong empirical evidence for SG rates of different species across ecosystems, and introduces a model (the SG model hereafter) that links sublinear growth with linear death and competition between species. In that model, the dynamics of the biomass of a species (*B*_*i*_) follows

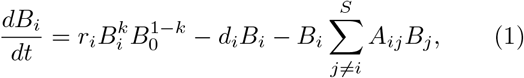

with *k* being smaller than 1 (*k ≈* 3*/*4). The model also includes a characteristic biomass *B*_0_ at which per-capita growth equals *r*_*i*_ (see Appendix), a linear death term (*−d*_*i*_*B*_*i*_) and the classical competitive term where the abundance of one species linearly impacts the growth of another (−*A*_*ij*_*B*_*j*_).

Before exploring the dynamics of the SG model, it is important to recall that stability and coexistence are two different properties (see Box). In the Generalized Lotka-Volterra (GLV) model for example, increasing species diversity leads to two consecutive results [7]. First, it disrupts community coexistence by pushing some species to extinction [5, 7]. Second, it destabilizes species abundances, leading to chaotic fluctuations and multiple stable states [6, 7, 15]. However, the two properties might not go hand-in-hand in other models: an increase in initial diversity could drive a community towards a more stable state (increased stability) where some species have nevertheless gone extinct (decreased coexistence).

In the present note we clarify how increasing the initial number of species affects stability and species coexistence in the SG model. First, we present a key particularity of the model, by which per-capita species growth diverges at low abundance, preventing species from going extinct (Section I). Second, we discuss the consequences of divergent growth on species coexistence (Section II). Third, we highlight that when divergent per-capita growth is reconciled with realistic assumptions, increasing diversity leads to competitive exclusion in the SG model and all species can no longer coexist (Section III).

## Box 1

**Diversity, stability and coexistence in ecological models**

**Diversity:** Diversity in ecology can be measured at different scales. In community ecology diversity typically refers to the number of different species in a community. Communities are often modeled as the result of an assembly process of species migrating from a regional pool [16]. The *final diversity* of a community is then usually lower than the *initial diversity* of the species pool due to species extinctions occurring during the assembly process.

**Stability:** Ecological stability encompasses various meanings and can be assessed through different measures [17–21]. Most often, stability refers to the capacity of an ecosystem to maintain its unperturbed state (resistance [22]) or go back to it (resilience [23]) following a perturbation. In theoretical works, community stability is often equated to linear stability. A community state is *linearly stable* if the real part of the dominant eigenvalue of its Jacobian is negative, so that abundances of all species will relax back into the steady state after a small perturbation. Stability can then be a mechanism of maintenance of species coexistence, by ensuring that a coexistence state is recovered after a perturbation [24, 25]. Stability, however, is not a direct proxy for species coexistence: for example, a state with no surviving species (trivial equilibrium) can nevertheless be stable. Also, loss of stability does not necessarily imply loss of coexistence: stability can be lost due to emerging fluctuations or exponential growth [26].

**Coexistence:** Two or more species are said to coexist if they persist at a positive abundance in the presence of the others. A main force at play against coexistence is *competitive exclusion*, which drives weaker competitive species to extinction [1, 3, 25]. A community state in which all of the initial *S* species coexist is often called *feasible* [27–29]. Coexistence does not require stability: species can persist together although not being at a steady state, as often observed empirical communities [26, 30].

It is noteworthy to remember that stability and coexistence are two different properties. The diversity-stability relation asks whether a system with more initial species becomes more or less stable to perturbations. The diversity-coexistence relation asks whether a system with more initial species has a smaller or larger probability to see species surviving together [3]. In the GLV model for example, the two relations go hand in hand: diversity disrupts both stability and coexistence [6]. In other models, however, the two mechanisms need not be equivalent: a system with more initial species can be more stable (positive diversity-stability trend) and at the same time drive species to extinction (negative diversity-coexistence trend). To explain the paradox behind the astounding levels of biodiversity in ecosystems we need to solve, at least, the second aspect: how can many species coexist without going extinct? [2, 3]

## I. DIVERGENT GROWTH AT LOW ABUNDANCES

A particularity of the SG model is that per-capita growth rates diverge at low species abundance. This means that when the number of individuals of a given species is low (due, for example, to competition), their growth rate becomes extremely large, thus automatically preventing the extinction of the species. To best see this, we can express the model in terms of population abundances *N*_*i*_ instead of the original biomass parametrisation (1). Dividing the total biomass of a species *B*_*i*_ by the typical individual biomass of each species *B*_0_ and rescaling the parameters accordingly, we get to (see Appendix)

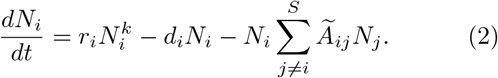

By dividing total species growth by the number of individuals, one obtains the per-capita growth rate

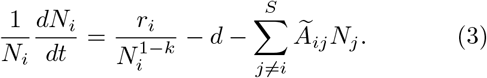

As species abundances becomes small, the per-capita perception of competition (the last term in the right hand side of equation 3) decreases linearly. Conversely, per-capita growth (the first term in the rhs) becomes infinitely large if *k <* 1 (Fig. 1). This leads us to the main ingredient of the SG model: Increasing the number of competitors pushes species towards lower abundances. At low abundance, however, competition becomes smaller while growth becomes extremely large. This has the effect of making the coexistence state divergingly stable, so that one could think that diversity favors species coexistence. In parallel, the states involving species extinctions are infinitely unstable, making species extinctions effectively unattainable (see Appendix).

**FIG. 1.**
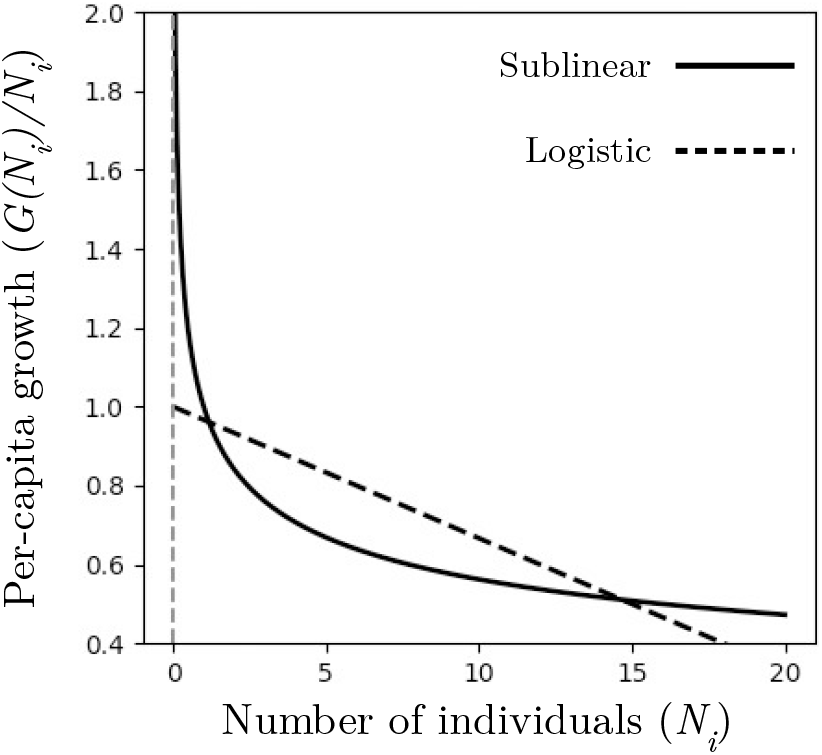
Per-capita growth diverges in the SG model. In the SG model, per-capita growth (dark line) diverges at low abundances. In the logistic model instead (dashed line), per-capita growth is finite at low abundance. Increasing the number of initial species in the SG model drives species to low abundances, where per-capita growth becomes extremely large. If growth diverges at low abundance, species can never go extinct due to competition.

If species cannot go extinct in the SG model, competitive exclusion is impossible by definition. Coexistence is then guaranteed by this unrealistic property. Furthermore, the lack of extinctions implies that the model cannot capture other fundamental effects in ecology, such as invasion-extinction processes nor community assembly: any new competitor that is added in a community will decrease the abundance of other species, but all species will remain in the community. Below we study the consequences of divergent growth.

## II. CONSEQUENCES OF DIVERGENT GROWTH ON SPECIES EXTINCTIONS AND COEXISTENCE

We study how many species survive (Fig. 2 left) and how many simulations end up in a stable state (Fig. 2 center) when the initial species diversity (*S*) increases (see Appendix). When all species survive at positive abundances and the final state is stable, the resulting system is said to harbor stable feasibility (Fig.2 right).

**FIG. 2.**
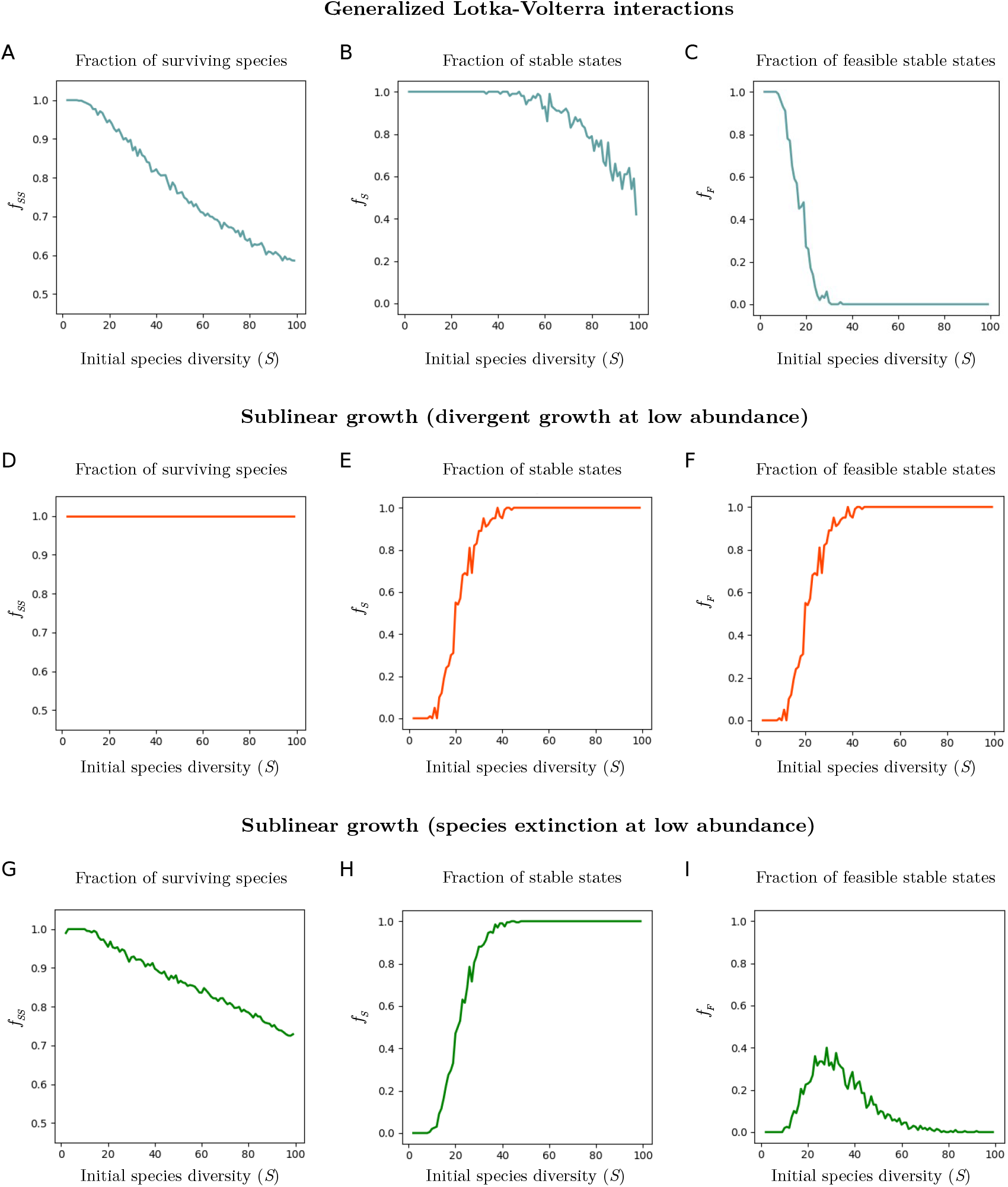
Extinctions, stability and stable feasibility under increasing initial diversity. Here we plot how the initial species diversity (*S*) modulates the fraction of surviving species at the end of simulations (left), the fraction of stable states (with negative dominant eigenvalue, center) and the probability that a community is both stable and all *S* species coexist (right). We plot the GLV model (A-C) and SG model (D-F) using the parameters of Figure 3A in [14], and the SG model with a minimal abundance of *B*_0_ = 0.03 below which species go extinct (G-I). The observation is that diversity increases stability in the SG model –a fundamentally new result to theoretical community ecology, but it has no effect on coexistence, because species can never go extinct. Yet if we allow species to go extinct, the SG model recovers a similar trend to the GLV model: increasing initial diversity leads to competitive exclusion, extinctions and loss of feasibility (see supplementary S5.2 in [14]).

Coexistence and stability follow the same trend in the GLV model [7]. Increasing initial diversity first leads to some species going extinct (Fig. 2A), and then leads to the remaining species losing stability (Fig. 2B, see Appendix). The combination of both processes implies that increasing species diversity decreases the likelihood of seeing a feasible stable state (Fig. 2C).

In the SG model instead, increasing initial diversity never drives species to extinctions, as competitive exclusion is artificially impaired by divergent growth. Coexistence is not affected by diversity (Fig. 2D). Conversely, increasing diversity stabilizes growth and fluctuations. In effect, for systems with a small number of initial species, species grow indefinitely. As diversity increases, competition can control growth dynamics and thus stabilizes the community (Fig. 2E, see Appendix). This new result is opposed to the negative diversity-stability trend of the GLV model.

The likelihood of finding stable feasibility increases with diversity in the SG model. Yet this is because stability increases, not coexistence (Fig. 2F, Fig. 3A in [14]). In the SG model, all species are always present at positive abundance, independently of the initial diversity or competitive strength of the community. Moreover, the SG model is proposed as an explanation for how loss of biodiversity could destabilize ecosystems [14]. However, if *S* is reduced, the model predicts a transition from stable dynamics towards chaotic or exponential growth. Because species do not go extinct, the SG model cannot predict the conditions for ecosystem degradation or secondary extinctions.

In sum, the main consequence of divergent per-capita growth rates is that species cannot go extinct. The key underlying mechanism is that increasing species diversity lowers the average species biomasses indefinetely, thereby stabilizing fluctuations (see Appendix and fig. 2E), with no impact on species extinctions (Fig. 2D). Below we unpack how diversity modulates species coexistence if species can effectively go extinct when their abundance falls below a certain threshold.

## III. RECONCILING THE MODEL WITH REALISTIC ASSUMPTIONS

We now introduce a more realistic behavior at low abundance, while keeping the empirical estimates of sublinear scaling in place (*k* = 3*/*4, see section S5.2 of the supplementary material of [14]). One way of doing that is to admit a biomass threshold *B*_0_ below which species go extinct [14]. One possible interpretation is that *B*_0_ is the typical biomass of a single individual, below which it is natural to assume species extinction.

In Figure 2 bottom, we plot the same analysis as previously but with an extinction threshold at *B*_0_ = 0.03 for all species. Under this realistic assumption, when diversity increases, species start to go extinct due to competitive exclusion (Fig. 2G). Species diversity and competition impair species coexistence in a more realistic SG model. Conversely, the trend by which diversity can stabilize dynamics remains in place (Fig. 2I), providing an interesting mechanism of community self-regulation. When coexistence of all species and stability are merged to measure stable feasibility, we can see that the SG model rarely finds stable survival of all the original species (Fig. 2I): either dynamics are unstable at low diversity, or species go extinct at high diversity.

The same result emerges from applying other mechanisms that limit the diverging growth rate in the SG model. This can be done e.g. by assuming that species growth rates are linear or constant instead of divergent below a certain threshold (see S5.2 of [14]). Additionally, the impossibility of extinctions on the SG model also depends on a key assumption on species competition: growth is sublinear with species abundances 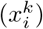, while the perceived impacts of competition are bilinear (*x*_*i*_*A*_*ij*_*x*_*j*_). As shown in [31], if interspecific interactions are also sublinear, one recovers the same results as in the GLV model (Fig. 2 top). Whether it is realistic to assume that intra- and interspecies effects scale differently with abundance remains to be tested in natural ecosystems (see e.g. [32, 33] for a discussion on evidence for sublinear interactions). In solid tumors, for example, one can consider that growth happens at the surface of the tumor where nutrients are available [34]. Resource competition with other cells or the attack of immune cells, which hardly penetrate the tumor, is also a surface process [35]. In that case, both growth and competition would be sublinear with tumor biomass and diversity would disrupt coexistence [31].

To conclude, increasing the realism of the SG model, e.g. by introducing a minimal biomass threshold for species survival, results in the same paradox of previous models: increasing the number of competing species decreases the likelihood of coexistence.

## IV. LOOKING AHEAD

In the present note we have discussed how diversity modulates stability species coexistence in the SG model. Our intention has been to unpack in simple terms certain results shown in [14].

On the on hand, diversity and competition can stabilize species growth in the SG model, uncovering a novel potential mechanism of community-level regulation of fluctuations [14, 31]. This is a new result for theoretical community ecology: it opens the door towards a more nuanced understanding of how species growth, interspecies interactions and community stability might be intertwined in natural ecosystems.

On the other hand, however, the absence of species extinctions in the SG model implies that competitive exclusion is impossible by definition. Hence, coexistence in the model emerges artificially. The SG model can be reconciled with species extinctions by assuming more realistic growth behaviors at low species abundances. Yet, when this is done, we recover the expected trend previously found in the GLV model: increasing species diversity leads to species extinctions and competitive exclusion.

Despite the interesting questions that the SG model raises, here we clarify the significant limitations it brings about and how to overcome them by allowing species to go extinct. Because species and ecosystems are increasingly endangered and their habitats and populations under threat, a fundamental task of ecological models is to understand and predict the conditions that can drive species to extinction.

## Appendix

### Typical biomass *B*_0_

The term *B*_0_ locates the typical biomass at which growth per unit of biomass equals the characteristic growth rate. If 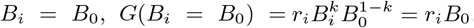, and in terms of growth per unit of biomass, one can write *G*(*B*_*i*_ = *B*_0_)*/B*_*i*_ = *r*_*i*_. If we define *B*_0_ to be the typical biomass of an individual, this term is then the per-capita growth rate.

This can be used to define a minimal biomass threshold: if the highest growth rate observed in a species is *r*_*i*_, then one would need that growth below *B*_0_ is no longer sublinear (see supplementary materials of [14], section S5.2). By imposing non-divergent growth or even extinction below *B*_0_, one recovers the condition by which excessive diversity disrupts species coexistence.

#### Rescaling from biomass to individuals

To rescale the model from biomass to number of individuals, one can assume a typical biomass of an individual of species *i* to be *B*_0,*i*_, which can be equivalent to the typical biomass at which growth is *r*_*i*_ (see above). Then, the number of individuals is *N*_*i*_ = *B*_*i*_*/B*_0,*i*_, and to rescale the equation we divide in both sides by *B*_0_,*i*

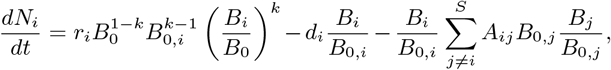

and one reaches equation (2) by rescaling competitive strengths by typical abundances Ã_*ij*_ = *A*_*ij*_*B*_*0*,*j*_ and assuming all species have the same typical biomass *B*_0,*i*_ = *B*_0_ *∀i*. In the GLV model, one should rescale 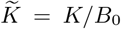 and again *Ã*_*ij*_ = *A*_*ij*_*B*_*0*,*j*_ to transform it into *dN*_*i*_*/dt*.

#### Parameter choices

The figures are based on replicating the studies in [14]. In figure 1, we use *r* = 1, *d* = 0 for the sublinear per-capita growth and *r* = 1 and 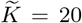 for the logistic per-capita growth. The parameters are taken only for illustrative purposes so that the two growth functions are comparable. In figure 2 we replicate the parameters used in Figure 3A of [14]: *r* = 1, *d* = 0, *k* = 3*/*4. We rescale logistic carrying capacity to *K* = 1, and the parameters of the interaction matrix are then sampled from a normal distribution with *µ* = 0.2 and *σ* = 0.1, equivalent to *µ* = 0.01 and *σ* = 0.005 in [14]. The expected diversity threshold of the GLV model is *S ≈* 128, computed from 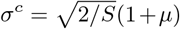 In figure 2 bottom we add a minimal biomass of *B*_0_ = 0.03, below which the abundance of a species is set to zero. Equivalent qualitative results as those of figure 2 are obtained when the threshold does not directly affect the abundance of a species, but only lowers its growth rate to avoid infinitely large values below *B*_*i*_ = *B*_0_ or *N*_*i*_ = 1 (not shown, but see supplementary section S5.2 of [14]).

#### Simulating the dynamics

Simulations in figure 2 are implemented by solving model (1) with a Runge-Kutta method of order 5(4) [36] via the solve ivp function of the scipy.integrate library, and we replicate the numerical methods described in [15, 37] to assess stability and surviving species. For each integer species value (x-axis), we generate 100 different interaction matrices *A*. For each, we generate random positive initial conditions with *x*_*i*_(*t* = 0) *∈* [0, 1] and run the dynamics for 1000 timesteps. After those, we compute the average number of species with strictly positive abundance over the 100 simulations (panel A), the fraction of simulations where all species abundances remain equivalent after 100 additional timesteps and the dominant eigenvalue of this state is negative (panel B) and the fraction of simulations in which the final state is stable and all species coexist at strictly positive abundance (panel C).

#### Linear stability and species extinctions

The link between divergent per-capita growth and the lack of species extinctions in the SG model can be understood by studying the Jacobian of the system. In the absence of an extinction threshold, the coexistence state where all *S* species survive(*B*_*i*_ *>* 0 *∀i*), has the Jacobian:

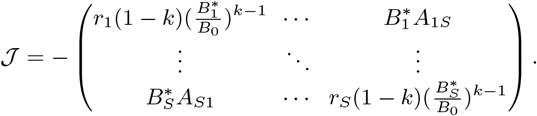

Increasing the number of species decreases their biomass due to competition. In that context, the off-diagonal terms emerging from interspecies competition become negligible compared to the diagonal terms, representing self-regulation, which diverge and push eigenvalues towards more and more negative values. This is how stronger competition can lead to a more stable state: increasing *S* makes the coexistence state divergingly stable.

Conversely, one can study the stability of the state with some species extinct (*B*_*i*_ = 0). In this state, the diagonal terms of the Jacobian are positive and infinite, making competitive exclusion unstable for any parameter setting. The same can be said by showing that any absent species can invade in the community: because growth rate diverges at low abundance, rare invaders will always survive and settle without driving other species to extinction.

## Acknowledgements

The authors thank O. Mazzarisi and the rest of the authors in [14] for positive and constructive discussions. G.A-G. thanks M. Barbier, Ll. Arola, B. Pichon, M. Lutterman and N. Humbert for insightful discussions and C. N. Adichie for inspiration. G.A-G. was supported by a 2022 postdoctoral fellowship of the Fundación Ramón Areces. J-F.A. was supported by the TULIP Laboratory of Excellence (ANR-10-LABX-41).

## References

1 G. Hardin, science 131, 1292 (1960).

2 G. E. Hutchinson, The American Naturalist 95, 137 (1961).

3 D. Tilman, Resource competition and community structure, 17 (Princeton university press, 1982).

4 R. M. May, Nature 238, 413 (1972).

5 G. Bunin, Physical Review E 95, 042414 (2017).

6 E. Mallmin, A. Traulsen, and S. De Monte, Proceedings of the National Academy of Sciences 121, e2312822121 (2024).

7 S. Marcus, A. M. Turner, and G. Bunin, arXiv preprint arXiv:2405.11360 (2024).

8 P. C. De Ruiter, A.-M. Neutel, and J. C. Moore, Science 269, 1257 (1995).

9 A.-M. Neutel, J. A. Heesterbeek, and P. C. De Ruiter, Science 296, 1120 (2002).

10 S. Johnson, V. Domínguez-García, L. Donetti, and M. A. Muñoz, Proceedings of the National Academy of Sciences 111, 17923 (2014).

11 E. Thébault and C. Fontaine, Science 329, 853 (2010).

12 J. Grilli, T. Rogers, and S. Allesina, Nature communications 7, 12031 (2016).

13 P. Landi, H. O. Minoarivelo, Å. Brännström, C. Hui, and U. Dieckmann, Population ecology 60, 319 (2018).

14 I. A. Hatton, O. Mazzarisi, A. Altieri, and M. Smerlak, Science 383, eadg8488 (2024).

15 G. Aguadé-Gorgorió and S. Kefi, Journal of Physics: Complexity (2024).

16 J. A. Drake, Journal of Theoretical Biology 147, 213 (1990).

17 S. Kéfi, V. Domínguez-García, I. Donohue, C. Fontaine, E. Thébault, and V. Dakos, Ecology letters 22, 1349 (2019).

18 V. Grimm and C. Wissel, Oecologia 109, 323 (1997).

19 I. Donohue, O. L. Petchey, J. M. Montoya, A. L. Jackson, L. McNally, M. Viana, K. Healy, M. Lurgi, N. E. O’Connor, and M. C. Emmerson, Ecology letters 16, 421 (2013).

20 V. Domínguez-García, V. Dakos, and S. Kéfi, Proceedings of the National Academy of Sciences 116, 25714 (2019).

21 F. Pennekamp, M. Pontarp, A. Tabi, F. Altermatt, R. Alther, Y. Choffat, E. A. Fronhofer, P. Ganesanandamoorthy, A. Garnier, J. I. Griffiths, et al., Nature 563, 109 (2018).

22 C. S. Holling, Engineering within ecological constraints 31, 32 (1996).

23 S. L. Pimm, Nature 307, 321 (1984).

24 E. Johnson and A. Hastings, arXiv preprint arXiv:2201.07926 (2022).

25 C.-Y. Chang, D. Bajić, J. C. Vila, S. Estrela, and A. Sanchez, Science 381, 343 (2023).

26 E. Benincà, J. Huisman, R. Heerkloss, K. D. Jöhnk, P. Branco, E. H. Van Nes, M. Scheffer, and S. P. Ellner, Nature 451, 822 (2008).

27 J. Grilli, M. Adorisio, S. Suweis, G. Barabas, J. R. Banavar, S. Allesina, and A. Maritan, Nature communications 8, 14389 (2017).

28 C. Song, R. P. Rohr, and S. Saavedra, Journal of Theoretical Biology 450, 30 (2018).

29 M. Dougoud, L. Vinckenbosch, R. P. Rohr, L.-F. Bersier, and C. Mazza, PLoS computational biology 14, e1005988 (2018).

30 C. Jacquet, C. Moritz, L. Morissette, P. Legagneux, F. Massol, P. Archambault, and D. Gravel, Nature communications 7, 12573 (2016).

31 O. Mazzarisi and M. Smerlak, arXiv preprint arXiv:2403.11014 (2024).

32 M. Barbier, L. Wojcik, and M. Loreau, Oikos 130, 553 (2021).

33 O. N. Mazzarisi, M. Barbier, and M. Smerlak, bioRxiv, 2024 (2024).

34 I. A. Rodriguez-Brenes, N. L. Komarova, and D. Wodarz, Trends in ecology & evolution 28, 597 (2013).

35 I.-K. Choi, R. Strauss, M. Richter, C.-O. Yun, and A. Lieber, Frontiers in oncology 3, 193 (2013).

36 J. R. Dormand and P. J. Prince, Journal of computational and applied mathematics 6, 19 (1980).

37 G. Aguadé-Gorgorió, J.-F. Arnoldi, M. Barbier, and S. Kéfi, Ecology Letters 27, e14413 (2024).

